# Understanding the Mechanisms Behind the Annuloplasty Effect in Tricuspid Valve TEER: A Computational Study

**DOI:** 10.1101/2025.11.04.686540

**Authors:** Collin E. Haese, Vijay Dubey, Mrudang Mathur, Felix Kreidel, Jan N. Fuhg, Issam Moussa, Tomasz A. Timek, Manuel K. Rausch

**Affiliations:** Department of Mechanical Engineering, The University of Texas at Austin, 204 E. Dean Keeton Street, Austin, TX 78712; Department of Cardiothoracic Surgery, Stanford University, 870 Quarry Rd Extension, Palo Alto, CA 94304; Clinic for Cardiology, Asklepios Hospital Harburg, Eißendorfer Pferdeweg 52, 21075 Hamburg, Germany; Department of Aerospace Engineering & Engineering Mechanics, The University of Texas at Austin, 2617 Wichita Street, Austin, TX 78712; Carle Heart and Vascular Institute, Carle Health, University of Illinois Urbana-Champaign Carle Illinois College of Medicine, 701 W Park St, Urbana, IL 61801; Division of Cardiothoracic Surgery, Corewell Health West, Michigan State University College of Human Medicine, 100 Michigan Ave SE, Grand Rapids, MI 49503; Department of Biomedical Engineering, The University of Texas at Austin, 107 W. Dean Keeton Street, Austin, TX 78712; The Oden Institute for Computational Engineering & Sciences, The University of Texas at Austin, 201 E. 24th Street, Austin, TX 78712

**Keywords:** Tricuspid Annulus, Remodeling, Annuloplasty, TriClip, Simulation

## Abstract

**Background:** An annuloplasty effect has been observed following tricuspid transcatheter edge-to-edge repair (TEER) and has been shown to have a therapeutic benefit. However, the mechanisms underlying the annuloplasty effect remain unknown.

**Objectives:** We investigate the impact of TEER-induced annular forces on the annuloplasty effect. Additionally, we explored the impact of clip size, clip orientation, leaflet pair, and leaflet site on TEER-induced annular forces.

**Methods:** We simulated 36 TEER repairs in finite element models of three human tricuspid valves. We used either a NTW or XTW TriClip. The clip was placed between either the anterior-septal or anterior-posterior leaflet pairs, in either the central or near-annulus site. For each scenario, we quantified the reduction in annular area, septal-lateral (SL) diameter, and anterior-posterior (AP) diameter. We also report the total annular force, orientation of maximum annular force, total papillary muscle force, leaflet stress, and coaptation area ratio following TEER.

**Results:** TEER induced annular forces, which strongly predicted the annuloplasty effect as measured by reduction in annular area and SL diameter. The maximum annular force aligned with the orientation of the clip. Furthermore, XTW clips induced more annular force and leaflet stress than NTW clips. A central anterior-posterior site induced more force than a near-annulus anterior-posterior site, but showed no difference from an anterior-septal pair.

**Conclusions:** TEER procedural parameters, such as clip size and site, strongly influence the magnitude of induced annular forces, which, in turn, correlate with the degree of annuloplasty effect.

## 1. INTRODUCTION

Tricuspid regurgitation (TR) is associated with an increased hospitalization rate, impaired quality of life, and overall poor prognosis^1^. Transcatheter edge-to-edge repair (TEER) has become a widely adopted percutaneous approach for treating patients with TR. This is, at least in part, due to its safety profile and ability to treat high-risk patient populations^2^. The TriClip (Abbott Structural Heart) and PASCAL (Edwards Lifesciences) devices are the two currently approved clip-based systems.

Interestingly, in addition to approximating the leaflets, TEER may also induce acute tricuspid annular remodeling. Thereby, it counteracts the annular dilation characteristic of secondary TR^3^. Several studies have reported on this TEER-induced reduction in annular dimensions, termed the “annuloplasty effect”^3–5^. Importantly, the annuloplasty effect is therapeutically important. Von Stein et al. found that patients with a larger reduction in annular area had superior survival over those with limited area reduction^3^. However, not all patients respond. For example, Russo et al. found that only two-thirds of patients experienced a reduction in annular diameter following TEER, while the remaining third of patients did not^5^. To date, no explanation has been identified for why the annuloplasty effect occurs only in some patients. This suggests a critical knowledge gap in our understanding of the mechanisms driving the annuloplasty effect.

Filling this knowledge gap may be a critical step in leveraging the annuloplasty effect to improve TEER outcomes. However, doing so is difficult in a clinical setting. For example, it is not possible to measure the forces that underlie the reduction in annular dimensions. Moreover, it is not possible to systematically vary clip positions and orientations in patients to compare outcomes. To overcome the limitation of clinical studies, in our current work we instead use our finite element models of the tricuspid valve. Therein, we have full control of TEER procedural parameters and can measure the resulting forces in the annulus, leaflets, and chordae. By implanting TEER devices of varying size and across varying positions in three high-fidelity virtual human tricuspid valve models, we explore the mechanisms behind the annuloplasty effect.

## 2. METHODS

### Healthy and Regurgitant Models

We used our previously developed high-fidelity, subject-specific finite element models of the human tricuspid valve to investigate the impact of TEER on valve mechanics and morphology. We used three valves (referred to as Valve #1, Valve #2, Valve #3) developed following the reverse engineering pipeline detailed by Mathur et al^6^. In brief, we previously mounted beating healthy human donor hearts in an organ preservation system to recreate a physiologic hemodynamic environment. For each heart, we recorded annular dynamics via sonomicrometry and transvalvular pressures throughout the cardiac cycle, which provided superior spatial and temporal resolution over echocardiography. Subsequently, we excised the tricuspid valve to characterize its geometry and material properties. We then combined these data to build a subject-specific finite element model of three healthy human tricuspid valves. To build a model with functional TR, we asymmetrically dilated the annulus and apically displaced the papillary muscles in each healthy valve model to induce a coaptation gap^7^. See Supplement A for more information on the healthy and regurgitant valve models.

### TEER Simulation

We simulated TEER on the tricuspid valves using Abaqus/Explicit 2024 (Dassault Systèmes). Clip geometries were based on digitized models of two TriClip sizes: the NTW and XTW (Abbott Structural Heart). Furthermore, we clipped all major cusps between either the anterior-septal (AS) or anterior-posterior (AP) leaflet pairs to reflect clinical practice^8^. Each pair was clipped either at a central site (Cent) or in a near-annulus site (NA). Both central and near-annulus sites were clipped for each leaflet pair when feasible. Only one clip was inserted in each simulation to exclude complex multi-clip interactions. The clip orientation was defined as the acute angle formed between the direction of the clip face and the anterior-posterior (AP) axis. The AP axis was defined for each valve as the line connecting the anteroseptal and anteroposterior commissures. In total, 36 repair cases were simulated with all procedural combinations summarized in Tables 1 and 2.

**Table 1.** Clip sizes used for simulating TEER in each valve, where n is the number of simulations.

| Clip Size | Valve #1 (n = 14) | Valve #2 (n = 10) | Valve #3 (n = 12) | Total (n = 36) |
| --- | --- | --- | --- | --- |
| NTW | 7 | 5 | 6 | 18 |
| XTW | 7 | 5 | 6 | 18 |

**Table 2.** Leaflet pairs and clip sites used for simulating TEER in each valve, where n is the number of simulations.

| Leaflet Pair and Clip Site | Valve #1 (n = 14) | Valve #2 (n = 10) | Valve #3 (n = 12) | Total (n = 36) |
| --- | --- | --- | --- | --- |
| AS NA | 2 | 2 | 2 | 6 |
| AS Cent | 4 | 2 | 4 | 10 |
| AP Cent | 6 | 2 | 4 | 12 |
| AP NA | 2 | 4 | 2 | 8 |

The TEER procedure was simulated in four steps as detailed in Figure 2. First, we applied a uniform pathologic transvalvular pressure of 42.5 mmHg to the ventricular side of the leaflets to bring the valve to its end-systolic configuration^9^. Next, we inserted an open clip with the valve still in its end-systolic configuration. Then, the transvalvular pressure was set to zero, and the valve returned to its end-diastolic configuration, which allowed the leaflets to settle onto the clip arms. We then closed the clip by bringing the arms together. Finally, we reapplied the transvalvular pressure and deployed the clip by releasing the fixed constraint at the clip base^10^. Simulation parameters and further details are provided in Supplement B.

**Figure 1.**
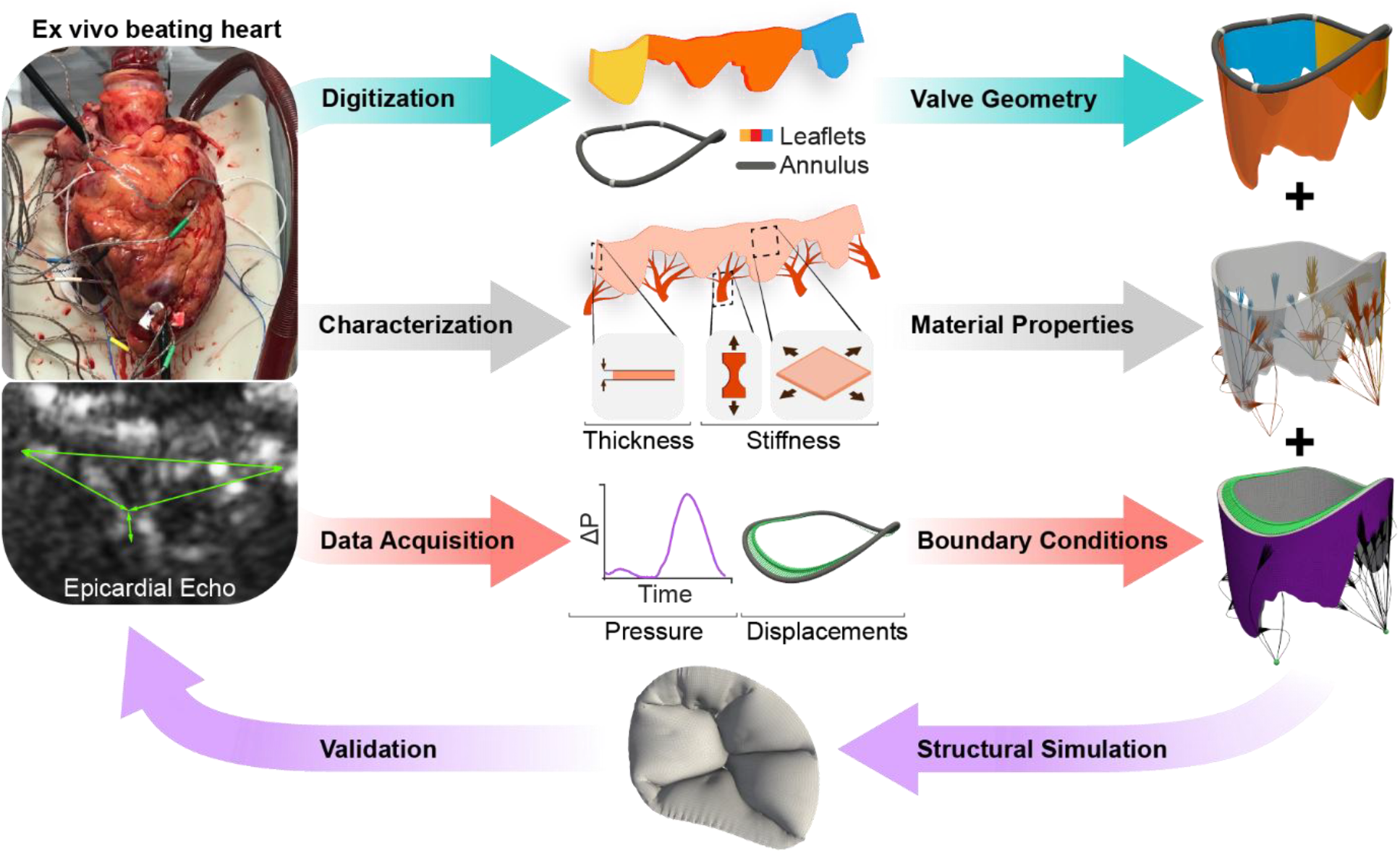
The reverse-engineering pipeline used to build patient-specific, high-fidelity finite-element models of the human tricuspid valves. Reproduced with permission from (Mathur et al., 2022).

**Figure 2.**
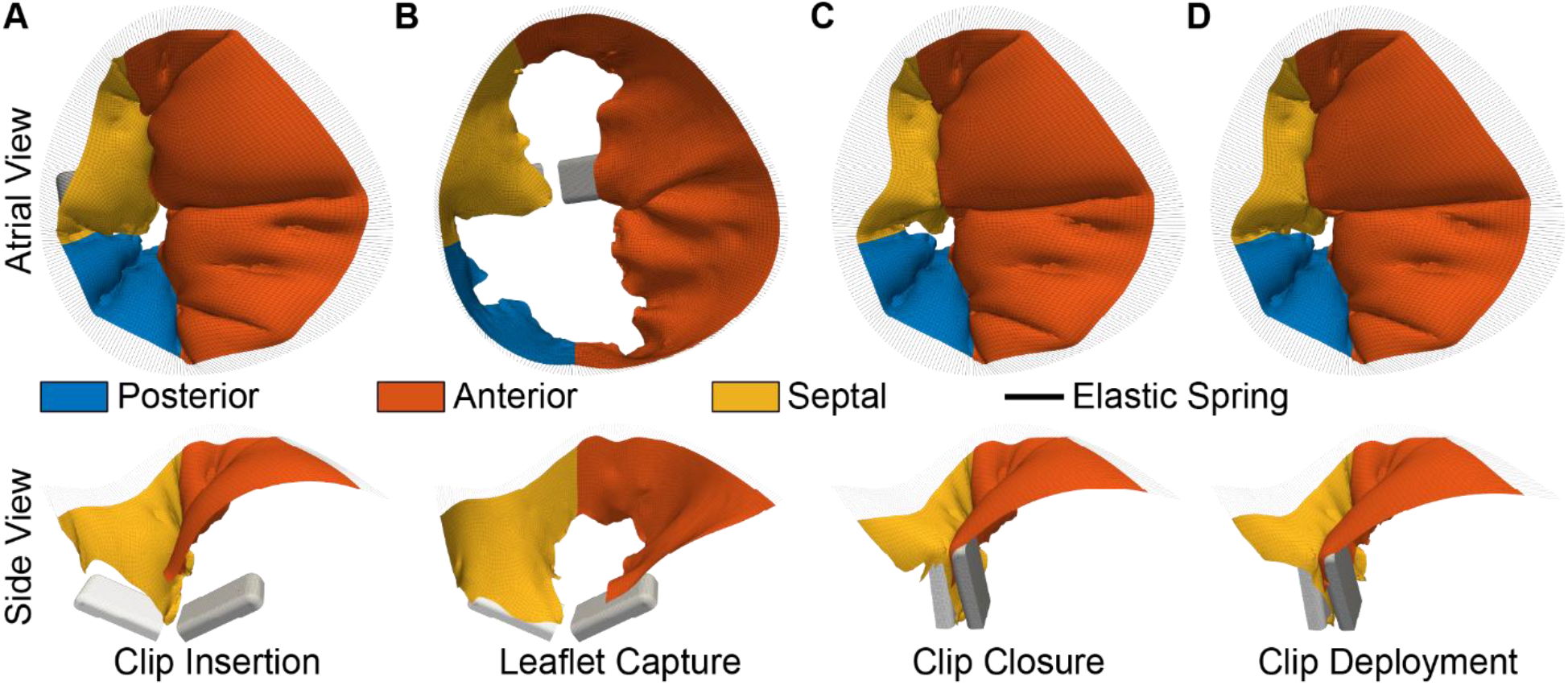
Example TEER simulation in the tricuspid valve, with the top row showing an atrial view, and the bottom row showing a cut view through the septal-lateral axis. **A**. The regurgitant valve model is pressurized to its end-systolic state, and the clip is inserted in an open configuration.**B**. We remove the transvalvular pressure and allow the leaflets to settle onto the clips. **C**. We close the clip and reapply the transvalvular pressure. **D**. We then deploy, i.e., release, the clip.

### Annular Boundary Conditions

In our previous work, we showed that annular boundary conditions affect predicted TEER outcomes^11^. Therefore, we accounted for the compliance of the perivalvular boundary by embedding the valve’s annulus into an elastic foundation of non-linear springs. The springs were calibrated to match the deformation observed in the healthy valves. This approach allowed for quantification of both the forces around the boundary as well as the amount of annuloplasty effect observed following TEER^11^. Further detail on the spring response is given in Supplement B.

### Repair Metrics and Statistical Methods

After simulating TEER, we quantified several measures of valve geometry, function, and mechanics and reported all as the difference between the pre- and post-clipped values. We report the reduction in annular area, septal-lateral (SL) diameter, and anterior-posterior (AP) diameter as geometric features. We report the change in total annular force, total papillary muscle (PM) force, leaflet stress, and coaptation area ratio as measures of valve mechanics. Supplement B describes in detail how each of these was computed. Additionally, we measured the orientation of the annular force by computing the acute angle of the axis connecting the two maximum spring forces on opposing leaflets with respect to the AP axis. We fit a linear regression to the change in annular force and each geometric feature to test their dependence. We used a linear mixed model to test the impact of clip size, clip site, and leaflet pair on annular forces using the *afex* library in R. Pairwise comparisons were performed using the *emmeans* library in *R*. Furthermore, we fit a linear regression to the maximum annular force orientation and clip orientation to test their dependence. We defined statistical significance as p<0.05.

## 3. RESULTS

### Annuloplasty Effect Correlates with Annular Forces

In our first step toward identifying the mechanisms of the annuloplasty effect, we quantified the annular forces induced by TEER. Moreover, we correlated those TEER-induced forces with the degree of the annuloplasty effect. The annuloplasty effect was measured as the reduction in annular area, SL, and AP diameter. Figure 3 shows area reduction as well as SL and AP diameter reductions against the total TEER-induced annular forces. We observed a clear and statistically strong correlation between forces and measures of the annuloplasty effect. Specifically, we found a coefficient of determination of R^2^=0.870 (p<0.0001) for reduction in annular area, R^2^=0.481 for SL diameter reduction (p<0.0001), and R^2^=0.106 for AP diameter reduction (p=0.0598). Absolute values for each are provided in Supplement Figure 7. In summary, TEER induces annular forces that in turn correlate with the annuloplasty effect. Interestingly, the annuloplasty effect appears to be driven primarily by a force-induced reduction in SL diameter and only secondarily by a reduction in AP diameter.

**Figure 3.**
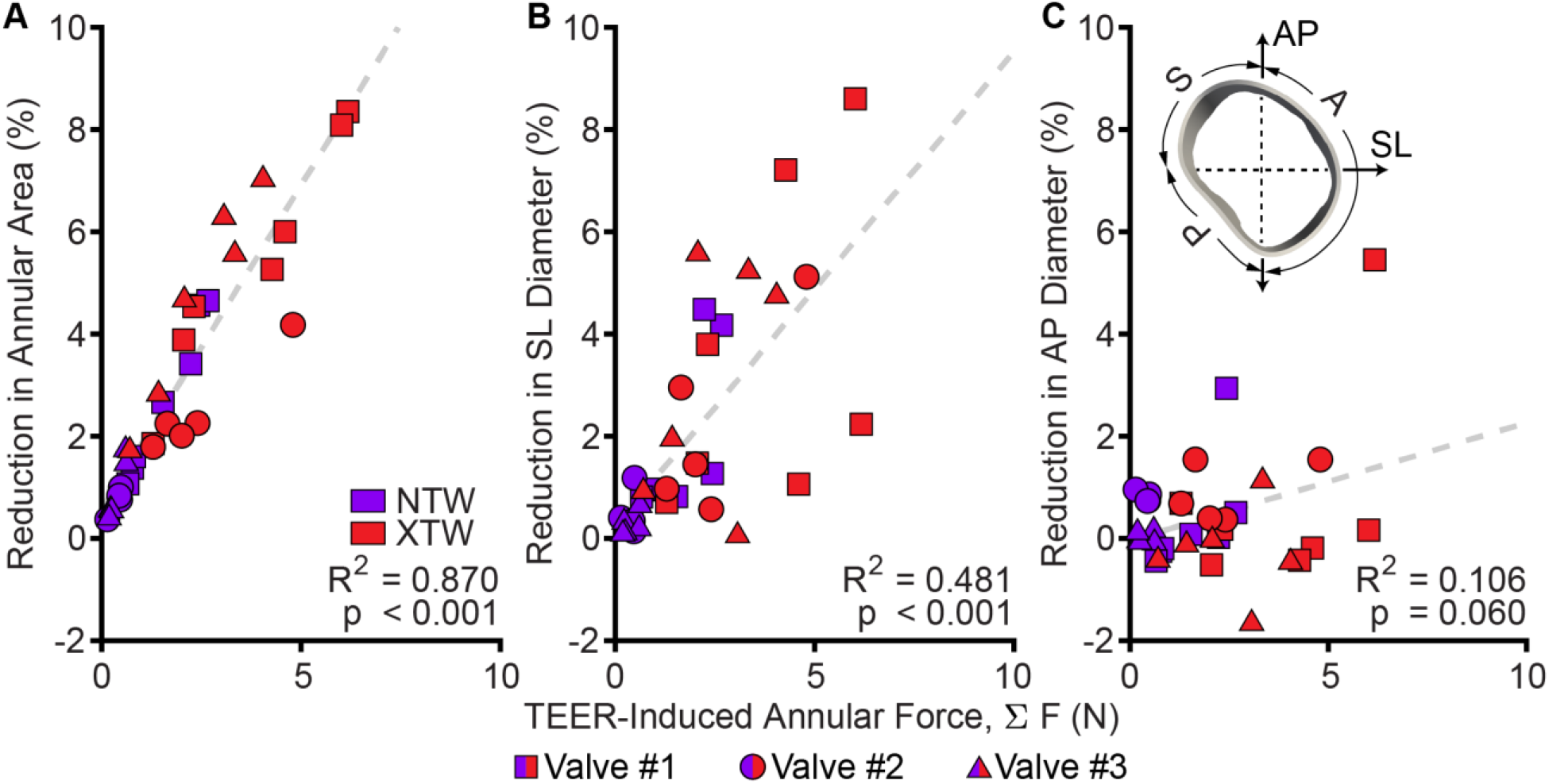
TEER-induced annular forces drive the annuloplasty effect. **(A)** Reduction in annular area and **(B)** septal-lateral (SL) diameter correlates with an increase in TEER-induced annular forces. **(C)** Annular forces and antero-posterior (AP) diameter reduction appear only weakly correlated.

### Annular Forces Depend on Clip Orientation

Given the important role of TEER-induced annular forces in driving the annuloplasty effect, we next investigated the distribution of those forces along the annulus. We found that annular forces were unequally distributed around the annulus. In fact, locations of peak forces correlated with clip orientation, as shown for three representative simulation results in Central Illustration A. To quantify the correlation between clip orientation and force distribution, we plot the annular force orientation against clip orientation in Central Illustration B. We found a strong correlation between both (R^2^=0.801, p<0.0001). In summary, clip orientation controls the distribution of annular forces. Notably, we did not find that the orientation of peak forces correlates with the annuloplasty effect (R^2^=0.030, p=0.330). In other words, in our models, clip orientation does not control annuloplasty effect, see Supplement Figure 8.

### Annular Forces Depend on Clip Size and Clip Site

In addition to exploring the role of clip orientation, we also varied clip size, clip site, and leaflet pair and quantified the resulting changes in TEER-induced annular forces. We found that the larger XTW clips induced larger forces than the smaller NTW clips (p<0.0001, 2.98±1.66 N vs. 0.91±0.84 N) and that a central clipping site – as opposed to a site near the annulus – also increased annular forces. However, the latter finding was only true when clipping the AP pair (p=0.0166, 2.59±2.05 N vs. 0.84±0.64 N, respectively), but not the AS pair (p=0.939). Interestingly, we found no difference in total annular forces when comparing the AS and the AP pairs (p=0.646). In summary, we find that larger clips increase annular forces and that clipping in a central site rather than near the annulus is advantageous when clipping the anterior and posterior leaflets.

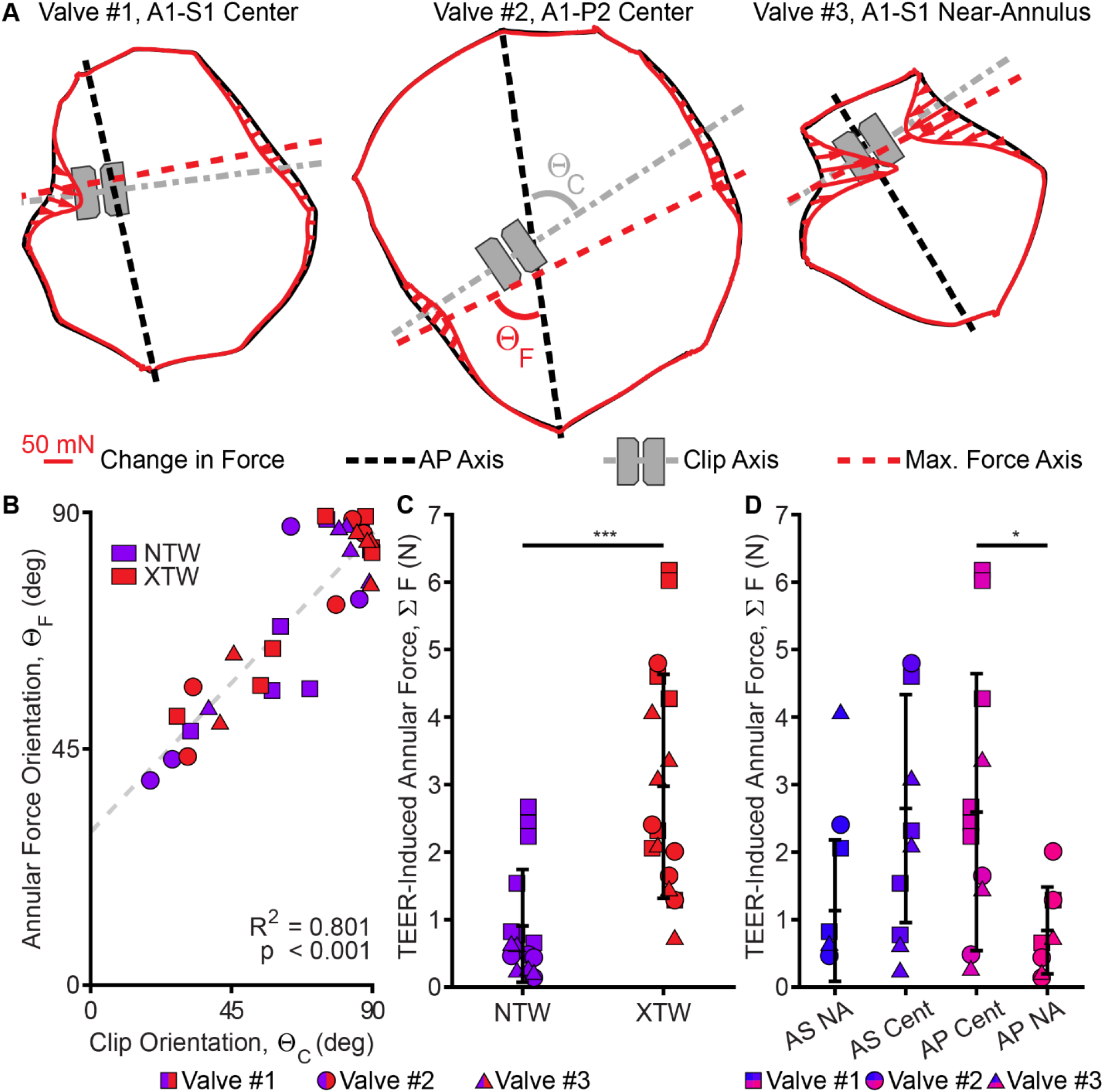
Central Illustration. TEER induces tricuspid valve annular forces. The largest induced annular forces align with the axis of the clip, and larger clips induce greater forces. **A**. Predicted post-TEER end-systolic configurations of three representative repair cases. The profile of the TEER-induced annular force is shown in red, along with the AP axis, clip axis, and maximum annular force axis. **B**. The orientation of the maximum induced annular force depends on the orientation of the clip, with both orientation angles defined with respect to the AP axis. **C**. TEER induces annular forces, and a larger clip induces more annular forces than a smaller clip. **D**. A clip located in a central AP site increases annular forces more than in a near-annulus AP site, but not significantly more than any AS site.

### Leaflet Stress Increases with Clip Size

Given that TEER-induced annular forces must be transmitted through the leaflets, we also explored the role of clip size, site, and leaflet pair on leaflet stress. In Figure 4A, we show the same valves as in Central Illustration A but overlaid with leaflet stress. We found that the larger XTW clips induced larger leaflet stresses than the NTW clips (p<0.0001, 62.80±48.99 kPa vs. 8.69±17.42 kPa), as shown in Figure 4B-C. Moreover, we found that stresses, just like annular forces, were larger when clipping the anterior and posterior leaflets centrally, rather than near the annulus (p=0.033, 53.34±52.64 kPa vs. 7.97±11.60 kPa, see Figure 4D). Here, too, we found that there are no differences in leaflet stress between the AP and AS pairs (p=0.961). In summary, leaflet stresses depend on clip size and site, but only marginally on leaflet pair.

**Figure 4.**
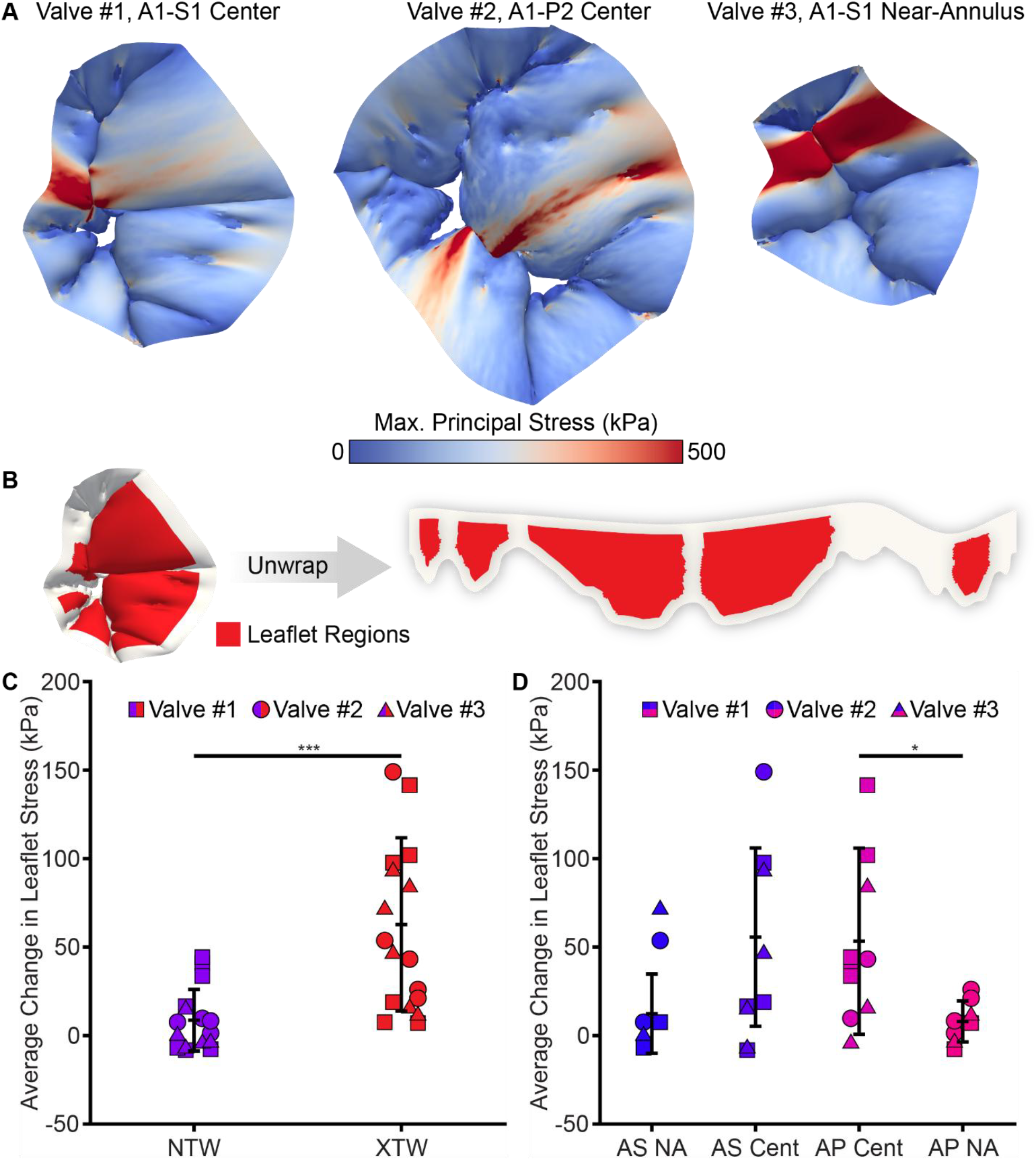
Larger clips induce more leaflet stress compared to smaller clips. **A**. Predicted post-TEER end-systolic configurations of three representative repair cases overlaid with contours of leaflet stress. **B**. The TEER-induced leaflet stress is computed element-wise across the central regions of each leaflet, shown in red. **C**. A larger clip increases leaflet stress, while a smaller clip does not substantially increase leaflet stress. **D**. A clip located in a central AP site increases stress more than in a near-annulus AP site, but not significantly more than an AS leaflet pair.

### Coaptation Area and Papillary Muscle Forces

Finally, we also computed TEER-induced changes in coaptation area and papillary muscle forces in response to clip size, site, and leaflet pair. Please find detailed results in Supplement Figures 9 and 10. In short, the coaptation area ratio increased following TEER (p=0.0312, 0.084±0.046), increased more with XTW clips than NTW clips (p<0.0001, 0.106±0.047 vs. 0.058±0.031), and was larger for the central AP site compared to a near-annulus AP site (p=0.0094, 0.107±0.049 vs. 0.047±0.018). For the TEER-induced total PM force, relative trends were similar to the other metrics. XTW clips significantly induced more PM force than NTW clips (p=0.0006, 1.16±1.22 N vs. 0.07±0.47 N). Additionally, a central AP clip site induced larger PM forces than either near-annulus site (p=0.0123, 1.36±1.24 N vs. 0.10±0.42 N anterior-posterior, p=0.0141, vs. 0.07±0.52 N anterior-septal).

## 4. DISCUSSION

In this study, we use our high-fidelity finite element models of human tricuspid valves to understand the mechanisms behind the TEER-induced annuloplasty effect. To this end, we varied clip sizes, clip sites, and leaflet pairs and computed the resulting annular forces, leaflet stresses, coaptation area, and papillary muscle forces. Moreover, we computed annular area reduction, as well as reductions in SL and AP diameters, as measures of the annuloplasty effect. In total, we simulated 36 repair cases in three different valves.

In summary, we found that TEER induces annular forces, which, in turn, strongly predict the annuloplasty effect as measured by reduction in annular area. Moreover, we found that the effect is primarily driven by a reduction in the SL diameter rather than the AP diameter. We also found that larger clips induced larger annular forces, increased leaflet stresses, and induced larger papillary muscle forces. Finally, we found that clips in the central versus the near-annulus site increased annular forces, leaflet stresses, and papillary muscle forces, but only when clipping the anterior and posterior leaflets, not when clipping the anterior and septal leaflets. Finally, we found that which leaflet pair is being clipped does not determine annular force magnitude, leaflet stresses, or papillary muscle forces.

Our findings mostly align with clinical observations and practice. First, our findings predict an annuloplasty effect after TEER, which has been observed in several studies^5,12^. Additionally, we find that SL diameter reduction drives the annuloplasty effect in our simulations, versus reduction in the AP diameter^4,13^. Lastly, our finding that larger clips yield largest annular forces and, thus, maximize the annuloplasty effect aligns well with the standard use of XTW devices^14^.

In contrast to clinical observations of TEER-induced annuloplasty, the mean amount of annular area reduction (3.0% vs 6.2%^3^) and SL diameter reduction (0.72 mm vs 2.5 mm^3^) was smaller in our simulations. However, this is a direct consequence of the chosen stiffness of the annular boundary, see Study Limitations. Surprisingly, we also found a limited dependence of TEER-induced annular force, and thereby annuloplasty, on which clip site and leaflet pair was chosen. Specifically, we found that the central clip site only increased annular forces compared to a near-annulus position within AP leaflet pairs. Annular forces between all other combinations of clip sites and leaflet pairs showed no significant differences. In comparison, Russo et al. found that clipping the AS pair had the greatest effect on reducing the annular diameter^5^. Others have also suggested that device positioning influences the annuloplasty effect^3,4^, which we failed to confirm.

### Translational Impact

There are several clinically relevant takeaways based on our models’ predictions. First, TEER induces an increase in annular forces. Larger TEER-induced annular forces strongly predict more annuloplasty. Widmann et al. suggested that TEER applies traction to the leaflets, which leads to annular remodeling, but lacked the means to directly measure this force^12^. To the best of our knowledge, this is the first study that investigates and quantifies the annular forces induced by TEER. Furthermore, our results show this increase in annular forces varies between patients and repair configurations, and understanding how TEER-induced forces impact outcomes may inform improved procedural strategies. Second, the maximum induced annular force follows the orientation of the clip. This suggests that interventions could leverage clip orientation to direct force along the most compliant segments of the perivalvular annulus and the right ventricular free wall. This could lead to a reduction in annular area and a decrease in regurgitation, which has been shown to have therapeutic benefits^3^. Third, larger clips induce more annular forces and recover more coaptation area. This finding supports the use of the newer, larger generation of clips over the smaller sizes when feasible. However, in addition to directing the forces through the leaflets to the annulus, we also observed that leaflet stresses increased along the clip axis. This is important as increased leaflet stresses are likely predictors of maladaptation, which may counteract repair benefits^15,16^. Thus, there is a potential trade-off between maximizing induced annular forces and recovering coaptation area while simultaneously increasing the stress on the leaflets.

### Study Limitations

This study is a computational investigation of the effect of TEER on the tricuspid valve. Each simulation obeys physical laws and provides detailed and accurate predictions of the acute post-repair valve configurations. However, our quantitative values in this study depend on subject-specific factors, including leaflet material properties and thicknesses. For example, the predicted reduction in annular area is a direct consequence of the stiffness of the spring-embedded annular boundary. While the stiffness chosen allows for realistic annular displacements^5,11^, it results in less mean annular area reduction than previously reported (3.0% vs 6.2%^3^). Therefore, we encourage the reader to focus on qualitative insight over quantitative insight. That is to say that comparisons within the models, i.e., for different clip sizes and positions, are qualitatively accurate, but specific numbers likely depend on patient specific factors. Future studies will be conducted using valve models built from echocardiographic images and validated against the post-procedural configuration. Lastly, longitudinal data from pre- and post-TEER timepoints are required to validate these models long-term predictive capability.

## 5. CONCLUSION

An increase in TEER-induced annular forces strongly predicts an annuloplasty effect. Larger clips and clipping in a central (versus near-annulus) location on anterior-posterior leaflet pairs predict larger TEER-induced forces, and therefore a larger annuloplasty effect. These also increase coaptation area and totally papillary muscle forces, while significantly increasing leaflet stresses which may trigger maladaptive responses. Additionally, clip orientation determines the location of peak annular forces. This suggests that interventions could direct force along the most compliant segments of the right ventricular free wall to maximize an annuloplasty effect. In conclusion, leveraging procedural parameters such as clip size, site, and orientation to maximize force on the annulus can increase the amount of TEER-induced annuloplasty.

## CLINICAL PERSPECTIVES

### What is Known?

TEER induces an annuloplasty effect in some patients but not all. Moreover, a larger annuloplasty effect has been associated with better outcomes.

### What is New?

Using three human-specific finite element models of the tricuspid valve, we identify the mechanisms of TEER-induced annuloplasty effect. Specifically, we find that the annuloplasty effect is a direct consequence of clip-induced annular forces, whose magnitude and orientation can be controlled through the judicious choice of procedural parameters.

### What is Next?

Using subject-specific, image-based models of patients’ tricuspid valves, our goal will be to provide recommendations of procedural parameters that maximize the annuloplasty effect and, thus, outcomes.

## ABBREVIATIONS

TEER: Transcatheter edge-to-edge repair
TR: Tricuspid regurgitation
PM: Papillary muscle(s)
AS: Anterior-septal leaflet pair
AP: Anterior-posterior leaflet pair
SL: Septal-lateral
Cent: Central clip site
NA: Near annulus clip site

## ACKNOWLEDGEMENTS

This research was supported by funding form the National Heart, Lung, and Blood Institute of the National Institutes of Health through award numbers 1F31HL178280 to C.E.H., 1R01HL165251 and 1R21HL161832 to M.K.R. and T.A.T. The authors acknowledge the Texas Advanced Computing Center (TACC) at The University of Texas at Austin for providing high-performance computing resources that have contributed to the research results reported within this paper.

