## SUPPLEMENTAL MATERIALS for "Understanding the Mechanisms Behind the Annuloplasty Effect in Tricuspid Valve TEER: A Computational Study"

### SUPPLEMENT A

#### Healthy and Regurgitant Valve Models

The full details of the reverse-engineering pipeline used to build the patient-specific finite element tricuspid valve models is given in the Texas TriValve 1.0 study by Mathur et al.[1] Each healthy valve model was built from a donated healthy human heart rejected from transplantation. Each heart was mounted, perfused, and paced in an organ preservation system. We measured annular motion, leaflet motion, and transvalvular pressure using sonomicrometry crystals, epicardial ultrasound, and pressure transducers, respectively, while paced to physiologic conditions. After recording the ex vivo behavior, we excised the valve and flattened the leaflets. We took images to record the shape and chordal insertion sites on the leaflets, and also measured the thickness of the leaflets and chordae tendineae. We recreated a three-dimensional valve geometry by nonrigidly transforming the leaflet geometry onto the ex vivo annular shape. We captured the mechanical behavior of each valve's leaflet using biaxial extension tests and fit these data to a Fung-type constitutive model. We captured the mechanical behavior of the chordae tendineae using uniaxial extension tests and fit these data to an incompressible Ogden material model. The ex vivo dynamics, valve geometry, and material data was combined to make a finite element model of each tricuspid valve. These were validated against the epicardial ultrasounds taken on the whole hearts. The total annular area and displacement of valve #2 was isotopically reduced by 5% to attain competent valve closure.

For each healthy valve, we modified the model to mirror the pathology of functional tricuspid regurgitation induced by pulmonary arterial hypertension. For each valve, the annulus was asymmetrically dilated to attain a 62% increase in the end-diastolic annular area and a circularity of one[2].The leaflet's free edge and the papillary muscle heads were passively dilated by the same amount. Furthermore, the papillary muscle heads were apically displaced by 5 mm. Leaflet and chordae thicknesses and material properties were held constant. The transvalvular pressure was set to 42.5 mmHg for each regurgitant valve[3]. These modifications induced coaptation gaps in each model, resulting in a model of functional tricuspid regurgitation for each valve.

### SUPPLEMENT B

#### Details on Finite Element Simulation of TEER

All simulations were conducted in Abaqus/Explicit 2024 (Dassault Systèmes, Vélizy-Villacoublay, France) on the Texas Advanced Computing Center's Lonestar6 system. Leaflets were meshed using linear quadrilateral S4R incompressible plane stress shell elements. Chordae tendineae were represented with multi-segmented three-dimensional linear T3D2 truss elements. For the clip geometry, the NTW clip arms were 6 mm wide and 9 mm long, and the XTW arms were 6 mm wide and 12 mm long[4]. Each pair of arms was spaced 1.4 mm apart[1,5,6]. Clips were meshed with rigid body R3D4 elements. For leaflet self-contact and contact excluding the clip face, general contact with a friction coefficient of 0.1 was used. Contact between the clip faces and the leaflets was assigned a frictional coefficient of 0.95. Contact was enforced with a linear pressure-overclosure behavior in Abaqus. All chordae tendineae were omitted from contact definitions. Damping forces due to the blood were approximated with a global linear bulk viscosity of 5.0 MPa·s.

We supported the valve annulus with non-linear springs and calibrated their stiffness to match the deformation observed in the healthy valves. The force-displacement response of the non-linear springs was given by $F = 0.01 * \exp(0.458 * u)$. The linear term of the force-displacement response was chosen by first calibrating linear spring to the healthy deformations. The springs' non-linear behavior penalized excessive annular deformations. We ultimately selected the stiffest calibrated response among the three valves to ensure consistency across all simulations. After clip deployment, unphysical clip motion was dampened by applying a viscous body force of 1e-6 $N/mm^3$ to the clip. A minimum time step of 5e-7 s was maintained with automatic uniform mass scaling. Additional details and background for these choices can be found in our previous studies using the Texas TriValve[1,5,7].

#### Details on Measures of Valve Geometry, Function, and Mechanics

All metrics were reported as the difference between the pre- and post-repaired values. The AP diameter was measured as the distance between the anterior-posterior and anterior-septal commissure. The SL diameter was measured as the distance perpendicular to the AP axis and passing through the septal-posterior commissure. Annular area was computed as the area of the annulus projected on to the 2D annular plane. We quantified the coaptation area ratio as the sum of the leaflet area engaged in contact at end-systole divided by the total leaflet area. The total annular force was calculated as the sum of the magnitudes of all spring forces. The total papillary muscle force was calculated as the sum of reaction force magnitudes at all papillary muscles. We reported leaflet stress as the element-wise difference in maximum principal Cauchy stress averaged over each major leaflet cusp.

### SUPPLEMENT C

#### Predicted Post-TEER End-Systolic Configurations

All predicted configurations are shown in Supplement Figure 1-6. The TEER-induced annular force profile and orientation for Valve #1 is shown in Supplement Figure 1, for Valve #2 in Supplement Figure 3, and for Valve #3 in Supplement Figure 5, respectively. Leaflet stress contours for Valve #1 are show in Supplement Figure 2, for Valve #2 in Supplement Figure 4, and for Valve #6 in Figure 6, respectively.

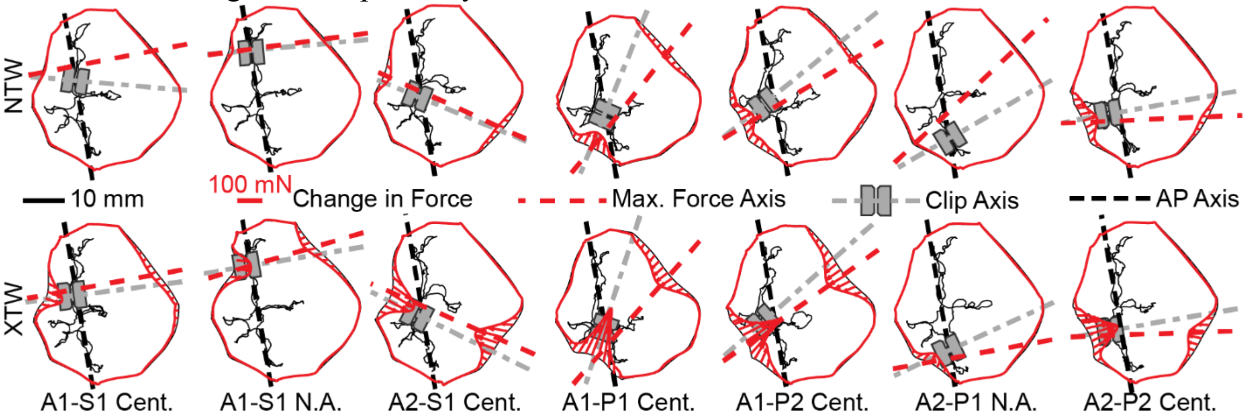


**Supplement Figure 1.** All predicted post-TEER end-systolic configurations for Valve #1. The profile of the TEER-induced annular forces is shown in red, along with the AP axis, clip axis, and maximum annular force axis.

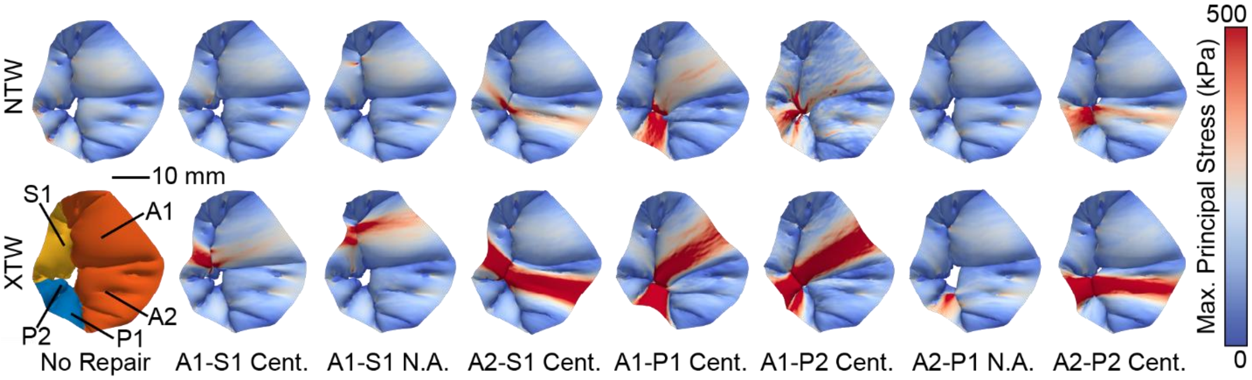


**Supplement Figure 2.** All predicted post-TEER end-systolic configurations for Valve #1 overlaid with contours of leaflet stress.

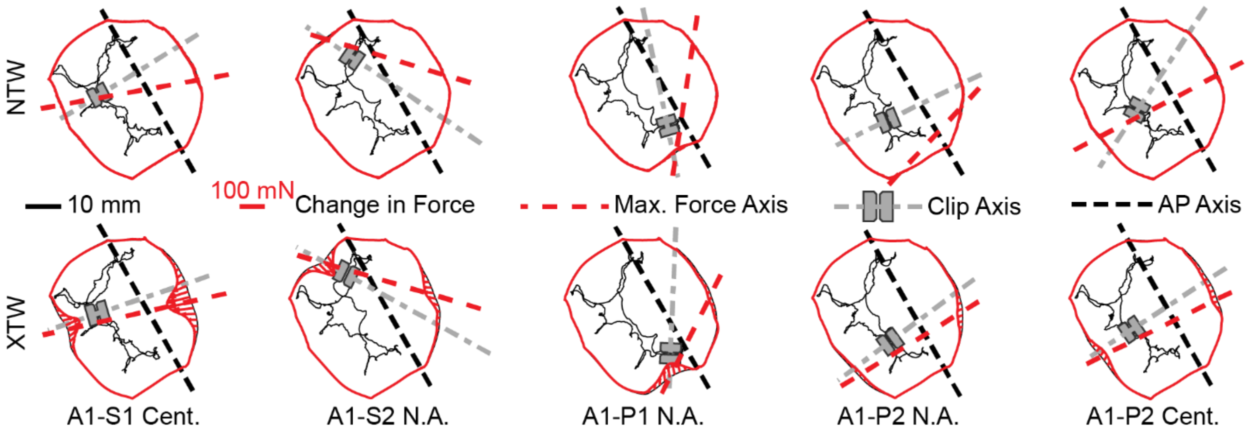


**Supplement Figure 3.** All predicted post-TEER end-systolic configurations for Valve #2. The profile of the TEER-induced annular forces is shown in red, along with the AP axis, clip axis, and maximum annular force axis.

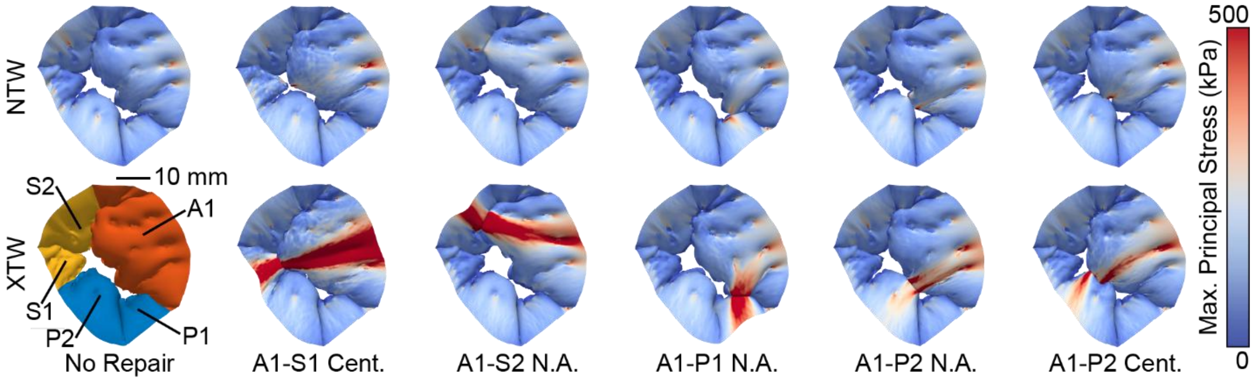


**Supplement Figure 4.** All predicted post-TEER end-systolic configurations for Valve #2 overlaid with contours of leaflet stress.

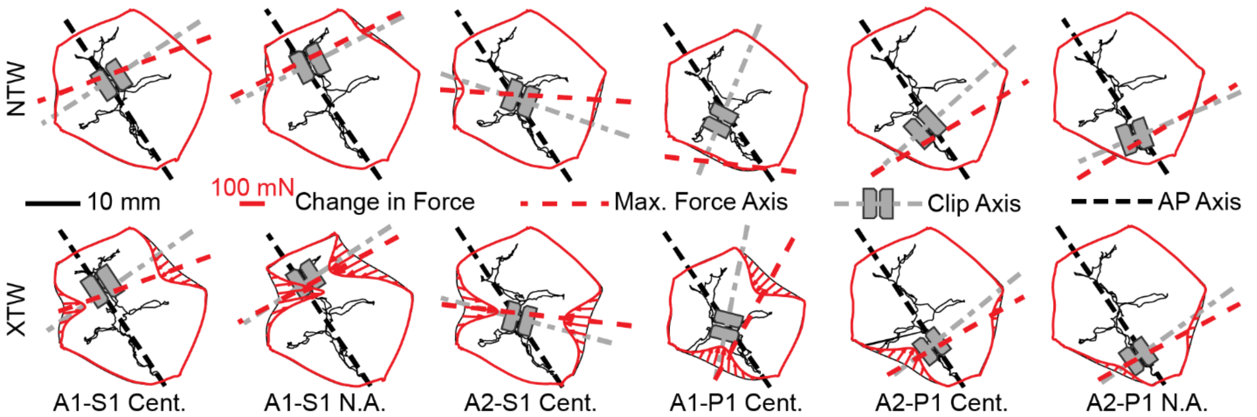


**Supplement Figure 5.** All predicted post-TEER end-systolic configurations for Valve #3. The profile of the TEER-induced annular forces is shown in red, along with the AP axis, clip axis, and maximum annular force axis.

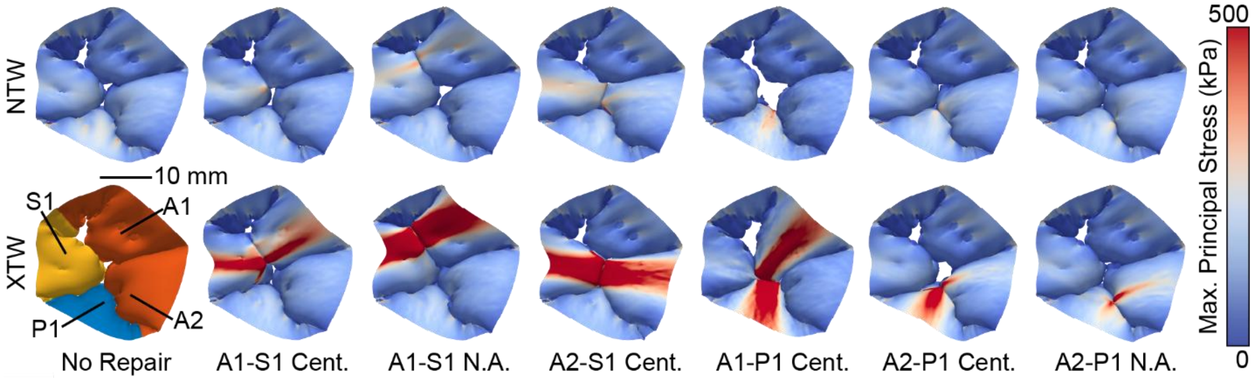


**Supplement Figure 6.** All predicted post-TEER end-systolic configurations for Valve #3 overlaid with contours of leaflet stress.

### Amount of Annuloplasty Effect Correlates with Annular Forces

We also measured the annuloplasty effect before converting to a percentage change. Supplement Figure 7 shows area reduction in $mm^2$ as well as SL and AP diameter reductions in mm against the total TEER-induced annular forces. Again, we see a clear and statistically strong correlation between forces and measures of an annuloplasty effect. Specifically, we find a $R^2$=0.946 ($p<0.0001$) for reduction in annular area, $R^2$=0.497 for SL diameter reduction ($p<0.0001$), and $R^2$=0.099 for AP diameter reduction ($p=0.056$). In summary, TEER induces annular forces that in turn correlate with the annuloplasty effect.

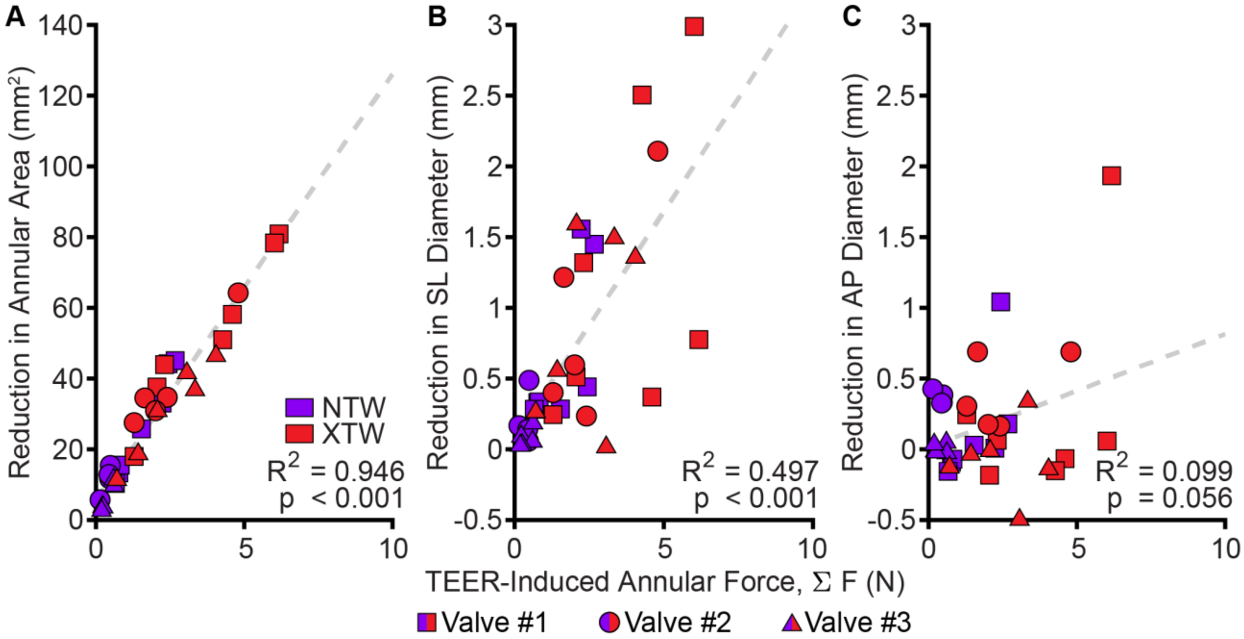


**Supplement Figure 7.** Measurements of the TEER-induced annuloplasty effect. **(A)** Reduction in annular area in $mm^2$ and **(B)** septal-lateral (SL) diameter in mm. **(C)** Reduction in antero-posterior (AP) diameter in mm.

#### Annuloplasty and Annular Forces Do Not Depend on Clip Orientation

To quantify the correlation between clip orientation and annuloplasty, we plot the reduction in annular area against clip orientation in Supplement Figure 8A. We also plot the total TEER-induced annular force against clip orientation in Supplement Figure 8B. We did not find a strong correlation between either ($R^2$=0.030, p=0.330, & $R^2$=0.041, p=0.248, respectively). In summary, clip orientation does not control annuloplasty effect nor the TEER-induced annular force magnitude.

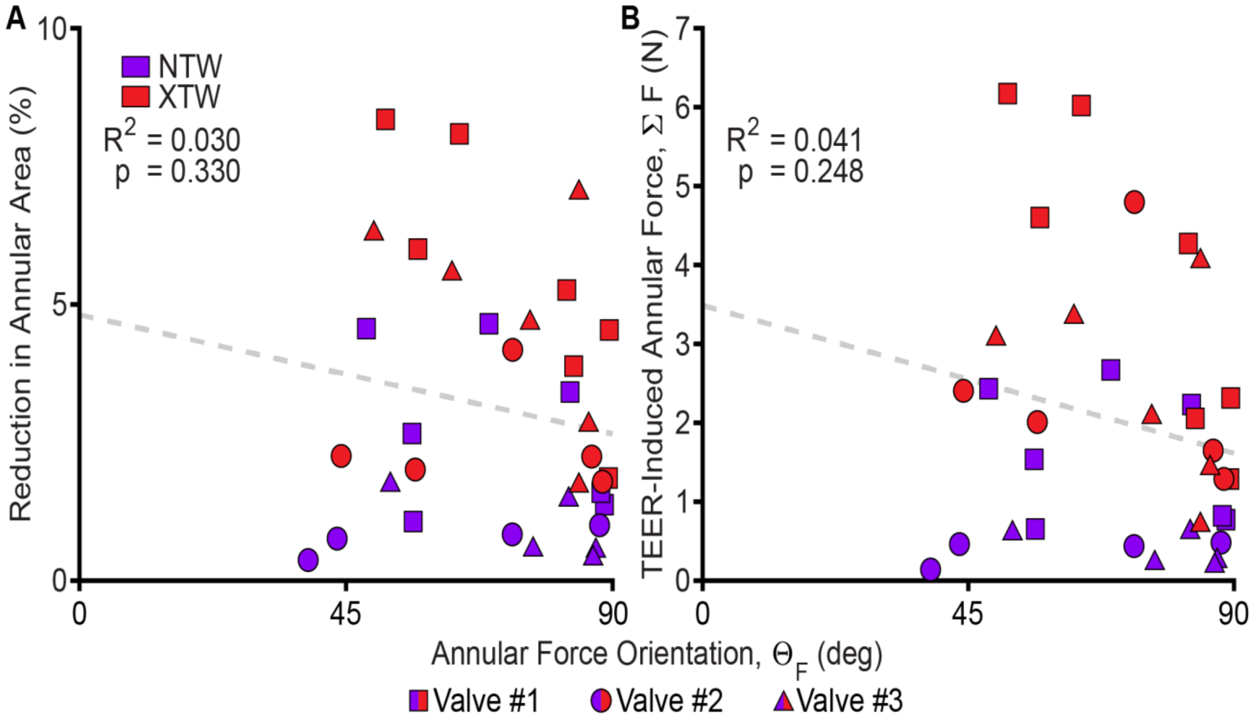


**Supplement Figure 8.** Clip orientation does not the control annuloplasty effect or induce forces. **A.** Reduction in annular area does not correlate with the orientation of the maximum annular force. **B.** Total TEER-induced annular forces do no correlate with the orientation of the maximum annular force.

#### Leaflet Coaptation Depends on Clip Size and Clip Site

Ultimately, TEER's primary function is to increase the coaptation area of the valve to reduce or eliminate regurgitation. Thus, we also explored the role of clip size, site, and leaflet pair on valve coaptation area. To quantify the amount of coaptation recovered following repair, we computed the difference in the coaptation area ratio, shown in Supplement Figure 9A, between pre- and post-TEER states. We found that coaptation area increased following TEER for all repairs (p=0.0312, 0.084±0.046). Supplement Figure 9B shows the change in coaptation area as a function of clip size. We found that the larger XTW clips recovered significantly more coaptation area than the smaller NTW clips ($p<0.0001$, 0.106±0.047 vs. 0.058±0.031). Supplement Figure 9C shows the change in coaptation area ratio as a function of clip site and leaflet pair. As with the annular forces and leaflet stresses, repair with a central AP clip site recovered more coaptation area than a near-annulus AP clip site (p=0.0094, 0.107±0.049 vs. 0.047±0.018). However, we find that there are no differences in change in coaptation area ratio between the AP and AS pairs (p=0.925). In summary, we find that the change in coaptation area ratio depends on clip size and site, but only marginally on leaflet pair.

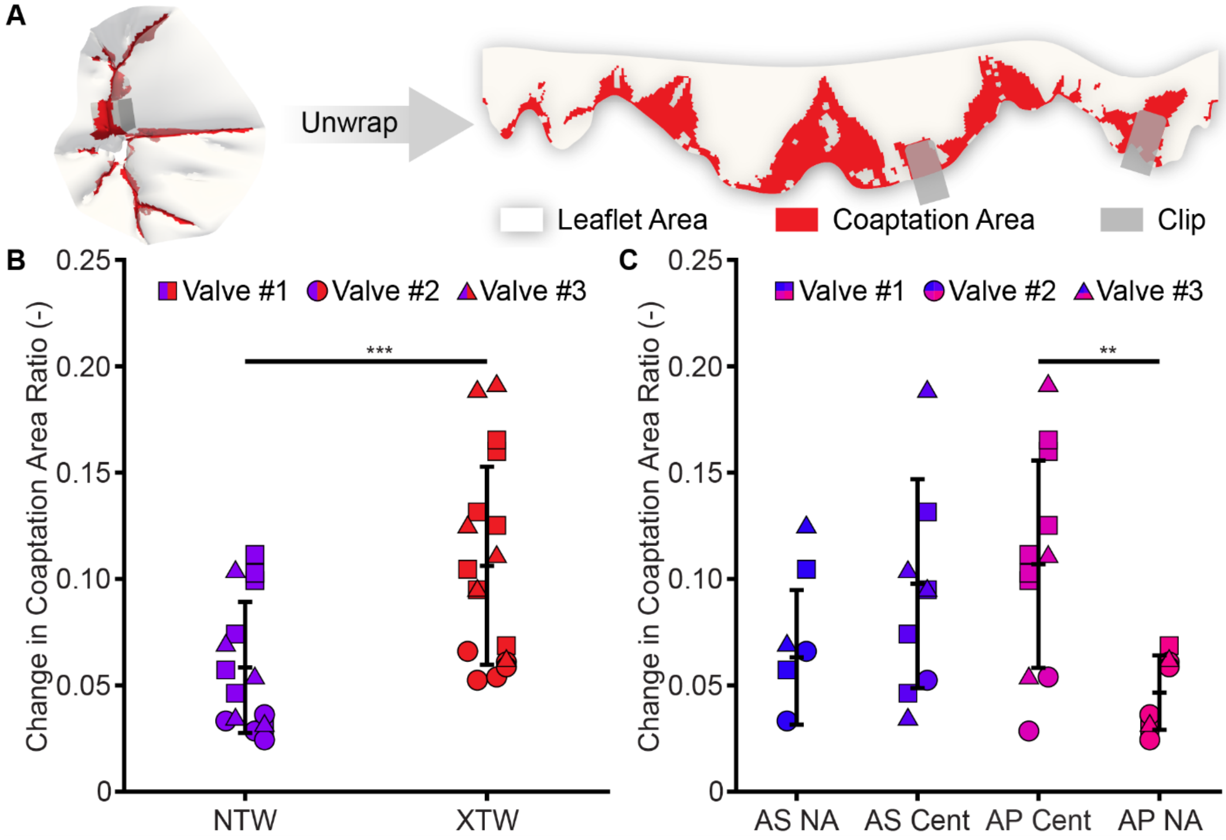


**Supplement Figure 9.** Larger clips recover more coaptation area. **A.** Coaptation area ratio is defined as the coaptation area divided by the total leaflet area. **B.** TEER recovers coaptation area, and a XTW clip recovers more coaptation area than an NTW clip. **C.** A central clip site between the AP leaflet pairs recovers more than in a near-annulus site, but not significantly more than in an either AS site.

### TEER-Induced Papillary Muscle Force Depends on Clip Size and Clip Site

In addition to investigating TEER-induced annular forces, we also investigated the impact of clip size, clip site, and leaflet pair on the total TEER-induced papillary muscle (PM) force. To quantify the TEER-induced PM force, we computed the difference in the sum of force magnitudes pre- and post-repair at each PM head as shown in Supplement Figure 10A. The TEER-induced PM force as a function of clip size is shown in Supplement Figure 10B. We found that XTW clips induced significantly greater PM forces compared to NTW clips (p=0.0006, 1.16±1.22 N vs. 0.07±0.47 N). Supplement Figure 10C shows the induced PM force as a function of clip site and leaflet pair. Repair with a central AP clip induced more PM forces than either near-annulus position, but there was no statistical difference between all other positions. We found that a central clipping site – as opposed to a site near the annulus – also increased annular forces (p=0.0123, 1.36±1.24 N vs. 0.10±0.42 N anterior-posterior, p=0.0141, vs. 0.07±0.52 N anterior-septal). However, there was no difference between clipping centrally in either leaflet pair (p=0.066, 1.36±1.24 N vs. 0.92±1.15

N). Additionally, we found no difference in total PM forces when comparing the AS and the AP pairs (p=0.095). In summary, we find that larger clips increase PM forces and that clipping in an AP central site rather than near the annulus increases PM forces.

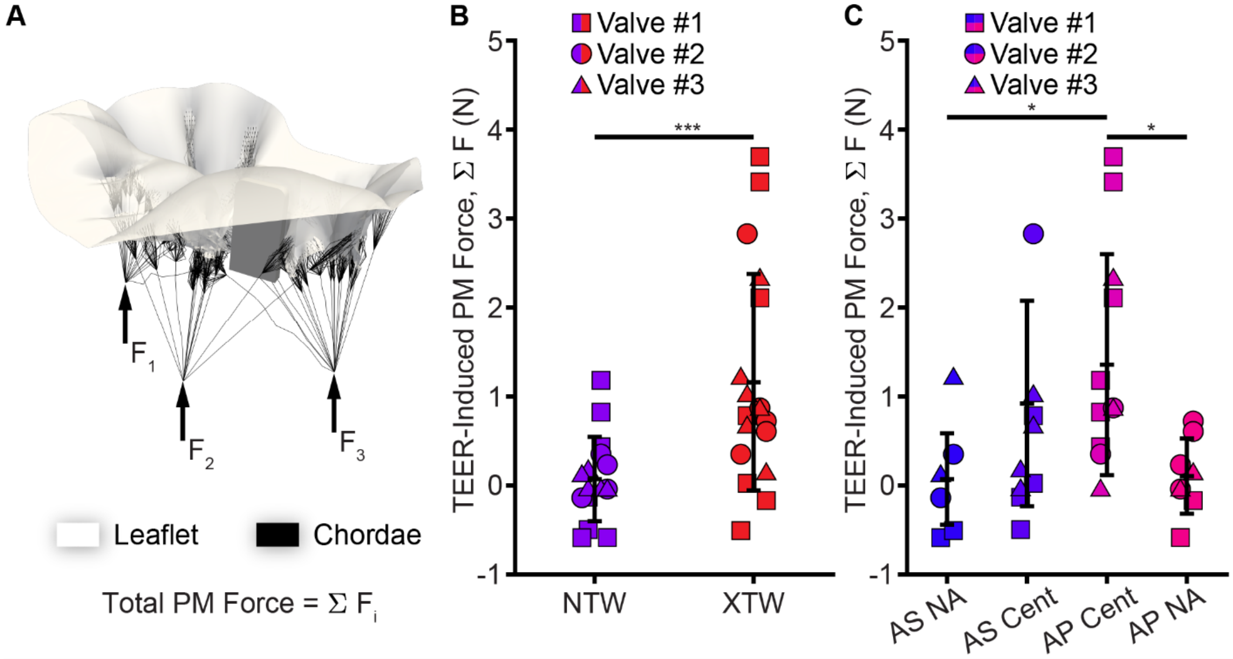


**Supplement Figure 10.** Total papillary muscle (PM) forces increase with larger clips. **A.** We compute the change in PM force as the difference pre- and post-repair in the sum of force magnitudes at each PM head. **B.** A XTW clip causes a larger increase in PM forces than an NTW clip. **C.** A central clip site between the AP leaflet pairs causes a larger increase in PM forces than clips located in a near-annulus position.

### SUPPLEMENTAL MATERIALS REFERENCES


1. Mathur M, Meador WD, Malinowski M, Jazwiec T, Timek TA, Rausch MK. Texas TriValve 1.0 : a reverse-engineered, open model of the human tricuspid valve. Engineering with Computers. 2022;38(5):3835-3848. doi:10.1007/s00366-022-01659-w

2. Ring L, Rana BS, Kydd A, Boyd J, Parker K, Rusk RA. Dynamics of the tricuspid valve annulus in normal and dilated right hearts: a three-dimensional transoesophageal echocardiography study. European Heart Journal - Cardiovascular Imaging. 2012;13(9):756-762. doi:10.1093/ehjci/jes040

3. Nickenig G, Kowalski M, Hausleiter J, et al. Transcatheter Treatment of Severe Tricuspid Regurgitation With the Edge-to-Edge MitraClip Technique. Circulation. 2017;135(19):1802-1814. doi:10.1161/CIRCULATIONAHA.116.024848
4. TriClip G4 Premarket Approval (PMA). U.S. Food and Drug Administration. https://www.accessdata.fda.gov/scripts/cdrh/cfdocs/cfpma/pma.cfm?id=P230007
5. Haese CE, Dubey V, Mathur M, Pouch AM, Timek TA, Rausch MK. Tricuspid valve edge-to-edge repair simulations are highly sensitive to annular boundary conditions. Journal of the Mechanical Behavior of Biomedical Materials. 2025;163:106879. doi:10.1016/j.jmbbm.2024.106879
6. Sturla F, Vismara R, Jaworek M, et al. *In vitro* and in silico approaches to quantify the effects of the Mitraclip® system on mitral valve function. *Journal of Biomechanics*. 2017;50:83-92. doi:10.1016/j.jbiomech.2016.11.013
7. Haese CE, Mathur M, Lin CY, Malinowski M, Timek TA, Rausch MK. Impact of tricuspid annuloplasty device shape and size on valve mechanics—a computational study. JTCVS Open. 2024;17:111-120. doi:10.1016/j.xjon.2023.11.002
